# Exploring Taxonomic and Functional Microbiome of Hawaiian Stream and Spring Irrigation Water Systems Using Illumina and Oxford Nanopore Sequencing Platforms

**DOI:** 10.1101/2022.07.01.498518

**Authors:** Diksha Klair, Shefali Dobhal, Amjad Ahmed, Zohaib Ul Hassan, Jensen Uyeda, Joshua Silva, Koon-Hui Wang, Seil Kim, Anne M. Alvarez, Mohammad Arif

## Abstract

Irrigation water is a potential source of contamination that carries plant and foodborne human pathogens and provides a niche for survival and proliferation of microbes in agricultural settings. This project investigated bacterial communities and their functions in the irrigation water from wetland taro farms on Oahu, Hawai’i using different DNA sequencing platforms. Irrigation water samples (stream, spring, and tank stored water) were collected from North, East, and West sides of Oahu and subjected to high quality DNA isolation, library preparation and sequencing of the V3-V4 region, full length 16S rRNA, and shotgun metagenome sequencing using Illumina iSeq100, Oxford Nanopore MinION and Illumina NovaSeq, respectively. Illumina reads provided the most comprehensive taxonomic classification at the phylum level where Proteobacteria was identified as the most abundant phyla in river stream source and associated wet taro field water samples. Cyanobacteria was also a dominant phylum from tank and spring water, whereas Bacteroidetes were most abundant in wetland taro fields irrigated with spring water. However, over 50% of the valid short amplicon reads remained unclassified and inconclusive at the species level. Whereas samples sequenced for full length 16S rRNA and shotgun metagenome, clearly illustrated that Oxford Nanopore MinION is a better choice to classify the microbes to the genus and species levels. In terms of functional analyses, only 12% of the genes were shared by two consortia. Total 95 antibiotic resistant genes (ARGs) were detected with variable relative abundance. Description of microbial communities and their functions are essential for the development of better water management strategies to produce safer fresh produce and to protect plant, animal, human and environmental health. This project identified analytical tools to study microbiome of irrigation water.

## INTRODUCTION

Irrigation water quality is a growing concern for agriculture as drainage is contaminated with agricultural runoff, wastewater overflows, and polluted storm or rainwater runoff, and irrigation waters are a potential source of plant and food-borne pathogens resulting in economic crop losses and human health risks (1, 2, 3). The microbial populations sharing the same niche may be commensal, symbiotic, or pathogenic. Many pathogenic bacteria can survive and proliferate in contaminated water and agricultural settings for long duration under favorable biotic and abiotic conditions (4, 5, 6). Studies have revealed that contaminated water splash can be a potential carrier of plant and food-borne pathogens (7, 8) that can enter plants through stomata, hydathodes and wounds (9). Also, antibiotics introduced through contaminated water are a continuing challenge as they may result in high selection pressure for antibiotic-resistant bacteria (10, 11, 12) and can persist even after water treatment.

Because of water scarcity and a simultaneous need to increase food production, there has been a shift from freshwater to alternative sources of irrigation water such as reclaimed or recycled water. However, potential health and environmental impact concerns are associated with the use of alternative water sources for irrigating the crops (13). Therefore, uncovering the bacterial composition and its associated functions in irrigation water will provide insight into formulating new disease management strategies and preventing major economic and public health risks. High-throughput sequencing has facilitated the identification of complex bacterial communities (14) independently of bacterial culture (15, 16). The bacterial microbiota is identified by analyzing the prokaryotic 16S ribosomal RNA (rRNA; ∼1,500 bp long) with nine variable regions interspaced between conserved regions. The 16S rRNA region selected for sequencing depends on the experimental objectives, design, and sample type. Sequencing of variable regions of the 16S rRNA gene using the most popular sequencing platforms, such as Illumina technology, uncovers the majority of bacterial microbiota (17). Illumina technology only permits sequencing of short variable regions of the 16S rRNA gene (18), and therefore, taxonomic assignment of reads at the species level may be elusive. Different species within a genus possess different phenotypic and virulence characteristics, therefore, accurate speciation of bacterial species is of utmost importance for formulating effective disease management strategies against pathogenic bacterial communities.

With the advancement in next generation sequencing technologies (NGS), 3^rd^ generation NGS technology, Oxford Nanopore enables generation of long sequence read lengths, possibly sequencing full length 16S rRNA genes (19). Full length sequences covering maximum nucleotide heterogeneity and discriminatory power allow better identification at the genus and species level. Comparative studies for Oxford Nanopore and Illumina 16S rRNA gene sequencing demonstrated similar bacterial composition at the genus level, although significant differences were observed at the species level (20). However, this technology complicates accurate species classification, particularly for bacterial species with a high sequence similarity in the 16S rRNA gene, owing to higher sequencing error rates (21).

Although Polymorphic marker gene (e.g., 16S rRNA, ITS) based analyses are useful for broad community taxonomical analysis, it did not provide functionality nor resolve the complexity of a microbiome. The shotgun metagenomic sequencing using advanced Illumina sequencing platforms have been proven to be a more reliable approach for these purposes (22). Metagenomic sequencing is a powerful tool for investigating occurrence, abundance, and distribution of ARGs in the natural environment and is suitable for discovery of novel ARGs that remain unidentified in culture-and amplicon-based analyses (23, 24).

This study aimed to investigate bacterial microbiota and associated gene function of different irrigation systems, mainly associated with wetland taro across the island of Oahu, Hawai’i. Mountain streams are the major source of irrigation waters used by farmers to irrigate crops. The overall goal of this project is to reveal the bacterial microbiota from different water source used for irrigation, in addition to field water, which is released back into the stream after use, carrying excess fertilizer, agricultural waste, ARGs and diverse unidentified bacteria. Bacterial communities were investigated based on 16S rRNA amplicon analysis using two principally different sequencing technologies and platforms—Illumina iSeq100 and Oxford Nanopore MinION and their taxonomic compositions were compared. The functionality of all the genes in complex samples and the distribution of ARGs were also investigated using shotgun metagenomic analyses. We aim to compare different technologies and approaches considered for microbiome studies such as shotgun metagenome, short-and long-amplicon read based to provide the desired level of accuracy in resolving the microbial taxonomic composition of the samples.

## MATERIALS AND METHODS

### Sample collection

Irrigation source and associated taro field water samples were collected in September - November 2020, across the Island of Oahu, Hawai’i. Irrigation water samples—R-S1-E, R-S2-W, R-S4-SE, and R-S5-SE—collected from natural streams which were sources of irrigation water for taro fields. Two water samples R-S7-N (stream emerging from the main reservoir on Oahu) and T-S6-N (tank storage water) were sources of irrigation for horticultural crops and other agricultural practices. Taro field water samples, R-F1-E, R-F2-W, R-F4-SE, and R-F5-SE, associated with R-S1-E, R-S2-W, R-S4-SE, and R-S5-SE, respectively, were collected to analyze bacterial microbiota. Two water samples, S-S3-N and S-F3-N were collected from a spring water source and an associated taro field, respectively. From each sampling site, 3 replicate water samples (2L per sample) were collected in sterile glass bottles, submerged 10 to 15 cm below the water surface. Samples were transported in an ice-cooler and processed in the laboratory for DNA isolation.

### Sample processing

Water samples collected from each site were vacuum filtered using the Millipore All-Glass Filter Holder kit (EMD Millipore Corporation, Billerica, MA). Collected water from each replicate was filtered through Whatman filter membrane to remove coarse to medium debris, followed by filtration through a MF-Millipore 8 µm sterile mixed cellulose ester (MCE) membrane (Merck Millipore Ltd., Tullagreen Carrigtwohill, Co. Cork, Ireland), and finally, filtered via MF-Millipore 0.22 µm sterile MCE membrane to trap the maximum bacterial community. The 0.22 µm membrane was used for bacterial DNA isolation using NucleoMag DNA/RNA Water Kit (MACHEREY-NAGEL Inc., Bethlehem, PA) following manufacturer’s instructions, with a few minor modifications to improve the DNA quantity and quality. The mechanical lysis was performed in lysis buffer MWA1 for 20 minutes using a vortex at full speed, followed by the addition of 25 µl of RNase (12mg/ml stock solution); the tubes were incubated for 15 minutes at room temperature (RT). A lysate of 450 µl was transferred to a 1.5 ml sterile Eppendorf tube and 25 µl of NucleoMag B-beads were added, mixed and shaken for 5 minutes, and kept on a magnetic rack at RT. The supernatant was removed, and the pellet was washed twice with buffer MWA3, followed by a single final wash with buffer MWA4. The magnetic beads were air dried for 15 minutes at RT; 70 µl RNase free water was used to elute DNA from the magnetic beads. Qubit dsDNA HS kit and Qubit 4 (Thermo Fisher Scientific, Waltham, MA) were used to quantify the genomic DNA. The DNA replicates from each sample were pooled for downstream processes and stored at -80°C.

### Illumina 16S rRNA library preparation, sequencing, and analysis

The polymerase chain reaction (PCR) was performed to amplify the V3-V4 hypervariable region of 16S rRNA gene following the reaction conditions: 94°C for 5 min; 40 cycles at 94°C for 20 s, 58°C for 30 s, and 72°C for 1 min; and the final extension at 72°C for 3 min. Primers 341F (5’-TCGTCGGCAGCGTCAGATGTGTATAAGAGACAGCCTACGGGNGGCWGCAG-3’) and 805R (5’-GTCTCGTGGGCTCGGAGATGTGTATAAGAGACAGGACTACHVGGGTATCTAATCC-3’) were used for PCR amplification (25). The amplified PCR amplicons were enzymatically cleaned using ExoSAP-IT (Affymetrix, Santa Clara, CA) and quantified using Qubit dsDNA HS Kit and Qubit 4. A secondary bead-linked transposome (BLT) PCR was performed using i5 and i7 adapters, provided in Nextera DNA Flex Library Prep Kit (Illumina, Inc., San Diego, CA), for barcode attachment (Supplemental Table 1). Each sample’s library was prepared in duplicate.

The BLT PCR conditions were initial denaturation at 98°C for 3 min, followed by X cycles of 98°C for 45 sec, 62°C for 30 sec, and 68°C for 2 min, with a final extension at 68°C for 1 min. The number of cycles of BLT PCR’s (X) was decided based on the amplicon concentration from the previous PCR as recommended by the manufacturer. Samples with concentrations ranging from 1-9 ng/µl and 9-21 ng/µl were subjected to 8 and 12 cycles BLT PCR, respectively. The amplicon libraries were cleaned using double-sided bead purification protocol following the manufacturer’s instructions. The purified libraries were quantified, normalized to 1 nM concentration and pooled. The pooled library was spiked with 2% using Phix control and loaded to Illumina iSeq100 for sequencing with a total of 302 run cycles to generate paired-end 150-bp reads. The total data yield was 717 MB with Q30 value of 88.1% and 89.6% for Read 1 and Read 2, respectively.

The sequenced data was base called and analyzed using BaseSpace sequence hub and EzBioCloud, respectively (26). The paired-end reads were used as a quality control to filter out low-quality (average quality value < 25) and merged using PandaSeq (27); primers were trimmed at a similarity cut-off of 0.8. The pipeline uses EzBioCloud database for taxonomic assignment and sequence similarity was calculated via pair-wise alignment. The chimeric reads with less than a 97% best hit similarity rate were removed using EzBioCloud non-chimeric 16S rRNA database through UCHIME (28). The sequenced data was clustered using CD-Hit7 and UCLUST with 97% similarity (29). Bacterial diversity was also analyzed and compared among the samples. For alpha diversity—OTUs, richness, and diversity were calculated, while for beta diversity—principal coordinate analysis (PCoA) and UPGMA clustering analyses were performed.

Valid reads were normalized for each sample to eliminate the bias produced because of variation in total number of reads. The Wilcoxon rank-sum test was used to calculate differences between the replicates. The differences in relative abundance in phyla and genera among the samples were determined using one-way ANOVA (single factor) with the least significant difference (LSD) test at α=0.05.

### Oxford Nanopore 16S rRNA library preparation, sequencing, and analysis

The genomic DNA of sample R-F1-E and S-F3-N was diluted to 1 ng/µl, and a total 10 µl gDNA was used for full-length 16S rRNA library preparation using 16S Barcoding Kit 1-24 (SQK-16S024; Oxford Nanopore Technologies, Oxford Science Park, UK) according to the manufacturer’s protocol. Ten µl of input DNA (10 ng) was mixed well with 25 µl LongAmp hot Start Taq 2X Master Mix and 5 µl of nuclease free water, afterward, 10 µl of each 16S barcode was added. The PCR was performed using following conditions: Initial denaturation at 95 °C for 1 min, 25 cycles of 95 °C for 20 sec, 55 °C for 30 sec and 65 °C for 2 min, with a final extension at 65 °C for 5 min. Each amplified sample was purified and washed with AMPure XP beads and 70% ethanol, respectively. For each sample, barcoded libraries were prepared in duplicate and quantified using Qubit Qubit 4; libraries were pooled to a desired ratio of 50-100 fmol in 10 µl of 10 mM Tris-HCl (pH 8.0) with 50 mM NaCl, and 1 µl of Rapid adapter (RAP) was added. The pooled library was loaded on to MinION vR9.4 flow cell and sequenced following manufacturer’s instruction. The generated sequencing data were monitored in real-time using the MinKNOW software (version 4.0.20). The obtained FAST5 files were base called using MinKNOW (version 4.0.20) embedded with Guppy version 3.2.10 pipeline. The generated full-length 16S rRNA sequence data were analyzed using cloud based EPI2ME (Oxford Nanopore) workflow for the identification of microbial community composition; EP2ME uses the NCBI GenBank database for taxonomic identification. The minimum and maximum read length of 1,500 and 1,600, respectively, were assigned as a quality control parameter, and Blastn was run using parameters max_target seqs=3 (finds the top three hits that are statistically significant) with blast e-value assigned as default 0.01. Per read coverage was calculated as the number of identical matches/query length. All classified reads were filtered for >77% accuracy and >30% coverage, which removed invalid alignments and were normalized for analysis. Results were obtained as comma-separated values (CSV) file via web report generated by EPI2ME workflow.

### Metagenomic library preparation, sequencing, and analysis

DNA from two samples (R-F1-E and S-F3-N) were used for preparing DNA metagenome libraries using NEBNext Ultra DNA Library Prep Kit (NEB, Ipswich, MA) following manufacturer’s instructions. The sonication-based method was used for fragmenting gDNA to the size of 350 bp. The obtained DNA fragments were end-polished, A-tailed, and ligated with full-length indexing adapters to the ends of the DNA fragments, followed by PCR amplification. The PCR products were purified using AMPure XP, and libraries were analyzed for size distribution and quantified using Agilent 2100 Bioanalyzer (Agilent, Santa Clara, CA) and real-time qPCR, respectively. The quantified libraries were pooled and sequenced on an Illumina NovaSeq 6000 platform to generate paired end reads. The obtained raw reads were pre-processed to trim low-quality bases with quality value (Q-value ≤ 38), reads with N nucleotides over 10 bp, and reads that overlapped with adapters over 15 bp. The obtained clean reads after quality control were assembled into scaftigs using MEGAHIT (30). The quality of the assembled data was predicted by N50 length. Scaftigs (≥500bp) were used for ORF (Open reading Frame) prediction using MetaGeneMark (31) and the ORF’s less than 100 nt were removed. Non-redundant gene catalogue, generated using CD-HIT (32), was further used to map clean reads using SoapAligner (33). Each metagenomic homolog was taxonomically annotated against NR database (34) for classification of microbial community at different taxonomic levels. For functional analysis, Kyoto Encyclopedia of Genes and Genomes (KEGG), evolutionary genealogy of genes: Non-Supervised Orthologous Groups (eggNOG), and Carbohydrate-Active enzymes (CAZy) databases were used for mapping functionally annotated unigenes. For Antibiotic Resistance Genes (AGRs) analysis, all the unique genes were BLASTp against the CARD (Comprehensive Antibiotic Research Database) database (*e*-value ≤ 1e^−5^). To identify the biologically relevant differences between two samples, statistical analyses were performed using STAMP v 2.1.3 (35), employing Fisher’s exact test with Newcombe-Wilson CI method (0.95 confidence interval) and Benjamini-Hochberg FDR correction factors and visualized using extended error bar plots.

## RESULTS

### Short length amplicon-based analysis—Illumina

The paired end 16S rRNA encoding gene sequences were obtained using Illumina iSeq100. After the data was pre-filtered and passed the quality check to remove low-quality, non-chimeric and non-target amplicons, the total number of valid reads with an average read length was computed (Supplemental Table 2) for each sample. Each sample was successfully sequenced in duplicate, except sample S-S3-N that encountered sequencing biasness in the 2^nd^ replicate run and failed to produce enough valid reads. After quality control, an average of 43,599 and 41,163 valid reads from the first and second replicate run, respectively, were obtained. In both the replicates, the highest and lowest number of valid reads were observed in sample R-S2-W (61,272 and 67,325) and R-F4-SE (22,274 and 26,908), respectively.

Based on phylum comparison performed using valid reads obtained from two sequencing replicates, no differences were observed, therefore the first replicate (barcode1-12) was considered for further taxonomic and diversity analysis (Supplementary Fig. 1). The valid reads generated from each sample were normalized to the least number of obtained valid reads (22,274; R-F4-SE) to overcome biasness in analysis outcomes. The reads were further clustered into operational taxonomic units (OTUs) at 97% identity ranging from 1,410 to 4,897. The OTU number remained higher in river stream sources, R-S1-E (3,416), R-S2-W (4,059), R-S4-SE (2,817), and R-S5-SE (4,897), compared with associated field water, R-F1-E (1,570), R-F2-W (2,753), R-F4-SE (1,978), and R-F5-SE (2,946). However, in spring source and field water samples, the OTU count remained comparable (Table 1). Furthermore, sample T-S6-N had the lowest count of 1,077 identified OTUs, followed by sample R-S7-N with 1,410 OTU numbers.

**Table 1.** List of total number of OTUs and calculated diversity analysis.

| Sample | OTUs | ACE | CHAO | Jackknife | Shannon | Simpson | Phylogenetic Diversity | Good's Coverage of Library (%) |
| --- | --- | --- | --- | --- | --- | --- | --- | --- |
| R-F1-E | 1,570 | 2,712.05 | 2,458.62 | 3,062.1 | 3.97 | 0.12 | 1,635 | 96.57 |
| R-S1-E | 3,416 | 5,924.43 | 5,273.62 | 6,074.57 | 5.35 | 0.08 | 4,165 | 92.35 |
| R-S2-W | 4,059 | 6,950.62 | 6,139.12 | 7,115.85 | 6.21 | 0.03 | 4,241 | 91.02 |
| R-F2-W | 2,753 | 3,513.98 | 3,231.47 | 3,649 | 5.17 | 0.08 | 3,230 | 95.98 |
| S-S3-N | 2,153 | 3,149.55 | 2,905.06 | 3,240.36 | 4.34 | 0.14 | 2,378 | 96.14 |
| S-F3-N | 2,157 | 3,526.22 | 3,221.37 | 3,746.8 | 4.35 | 0.1 | 1,435 | 95.49 |
| R-S4-SE | 2,817 | 4,123.62 | 3,730.12 | 3,994.1 | 5.28 | 0.04 | 3,516 | 94.86 |
| R-F4-SE | 1,978 | 2,408.57 | 2,214.96 | 2,521 | 4.42 | 0.11 | 2,473 | 97.56 |
| R-S5-SE | 4,897 | 7,684.84 | 6,782.04 | 7,219.57 | 6.86 | 0.01 | 4,739 | 89.97 |
| R-F5-SE | 2,946 | 4,257.84 | 3,804.92 | 4,166.01 | 4.91 | 0.09 | 3,116 | 94.56 |
| T-S6-N | 1,077 | 1,656.5 | 1,522.44 | 1,763.94 | 2.88 | 0.34 | 1,106 | 97.93 |
| R-S7-N | 1,410 | 1,818.37 | 1,697.27 | 1,859.62 | 4.97 | 0.02 | 1,359 | 97.99 |

### Taxonomic classification at phylum, genus, and species levels

Based on Good’s coverage index, the sequencing covered more than 94% of the taxonomic richness except for sample R-S1-E (92.35%), R-S2-W (91.02%) and R-S5-SE (89.97%; Table 1). A total of 18 phyla with relative abundance of >1% were compared after being identified in at least one sample (Figure 1A). Proteobacteria, a phylum with major plant and food-borne pathogens, was significantly the most abundant phylum in 12 different samples (Supplemental Table 3). The relative abundance of Proteobacteria was higher in river stream source samples, R-S1-E (76.99%), R-S2-W (71.28%), R-S4-SE (83.71%), and R-S5-SE (52.04%), and associated field samples, R-F1-E (66.57%), R-F2-W (78.64%), R-F4-SE (89.08%), and R-F5-SE (75.70%). Considering samples collected from North Oahu, Cyanobacteria was the topmost abundant phylum identified from the spring water sample S-S3-N (35.86%) and stored tank water sample T-S6-N (58.39%).

**Figure 1.**
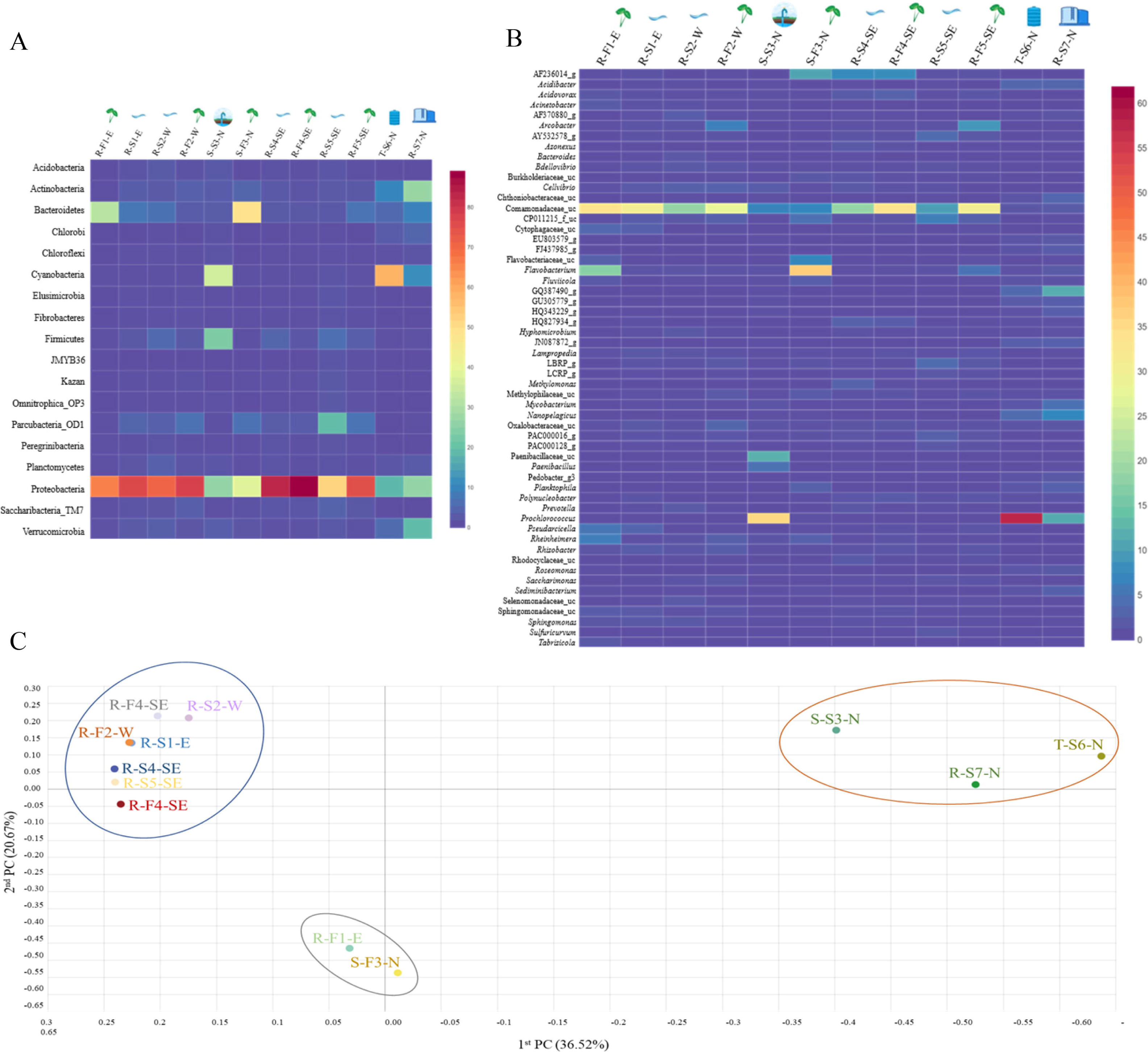
The distribution heatmap of bacterial **A)** phylum and **B)** genus detected with relative abundance >1% among all the water samples sequenced using Illumina iSeq100, an amplicon sequencing platform and analyzed on EzBioCloud. The heatmap was generated using displayR. **C)** Principal Coordinate Analysis (PCoA) clustering based on Bray-Curtis dissimilarity index was analyzed at genus level bacterial structure to visualize the variation in bacterial community structures among 12 different samples, forming three distinctive clusters. Cluster 1 (blue circle) shows close microbial communities of river streams and associated field samples, irrespective of geographical location. Cluster 2 (red circle) represents close microbial association between samples collected from North Oahu. Cluster 3 (gray circle) shows close microbial association between sample R-F1-E and S-F3-N.

Bacteroidetes was the most dominant phylum in spring water irrigated field with relative abundance of 48.82% and interestingly this phylum was also higher in the stream water irrigated field sample, R-F1-E (31.63%), whereas it remained <6.9% of relative abundance in other river stream source and associated field water samples. Phylum Actinobacteria was relatively higher in the reservoir stream source, R-S7-N (26.82%) compared with other samples. Other identified phyla varied in their relative abundance among all the samples, as shown in Figure 1A. The normalized valid reads from all the 12 samples were classified and compared at the genus level (Figure 1B). The taxonomic classifier used to classify valid reads identified uncultured genera and best hit genera classified with high and low confidence values, while the rest remained unclassified at a taxonomic level (genus-species). The genera within the family Comamonadaceae were classified as significantly most abundant among all the other identified genera and named as Comamonadaceae_uc by the taxonomic classifier (Supplemental Table 4). The taxonomic classifier could not differentiate the genera within the family Comamonadaceae owing to low confidence value in assigning the best hit to the reference database—indicating that short amplicon reads might not be efficient in classifying valid reads with high accuracy. The abundance of Comamonadaceae_uc was relatively higher in natural stream sources and associated field samples. *Prochlorococcus* was the most abundant genus identified in samples T-S6-N (58.3%) and S-S3-N (35.65%) collected from North Oahu. Spring field water sample S-F3-N was dominated by the genus *Flavobacterium* with relative abundance of 37.14%, while 16.99% *Flavobacterium* abundance was calculated in sample R-F1-E—the abundance remained <1% in all the other river stream and associated field water samples. The classified reads at the genus level, with a relative abundance of <1%, ranged between 22.32 - 61.87% among all samples, indicating diverse microbiota associated with different samples. The percentage of valid reads that remained unclassified varied between 4.83% (T-S6-N) and 21.55% (R-S4-SE) among all the samples (Supplemental Table 5).

At the species level, valid reads that remained unclassified among all the 12 samples ranged from 11.2% (T-S6-N) to 62.23% (R-F4-SE) (Supplemental Table 5). A total of 34 species classified at species level using EzBioCloud with relative abundance of more than 1%, only three species, *Flavobacterium fontis*, *F. hydatis* and *F. shanxiense,* remained classified with a high confidence value—indicating that the short length reads-based approach for classifying at species level is an inadequate approach for attaining species level resolution (Supplementary Fig. 2).

### Alpha and Beta diversity analyses

Non-parametric analysis of diversity indices, such as ACE, CHAO, and Jackknife, indicated higher bacterial diversity in river stream compared to associated field water samples, followed by sample S-F3-N, S-S3-N, R-S7-N, and T-S6-N (Table 1). The higher Shannon diversity indices of river stream source field water indicated an increased abundance and bacterial community than associated field water; however, a negligible difference between spring source S-S3-N (4.34) and field water S-F3-N (4.35) was observed (Table 1). The Shannon diversity calculated for sample T-S6-N and R-S7-N was 2.88 and 4.97, respectively.

Taken together, natural stream source water contaminated with fertilizer runoff, wastewater runoff and other agricultural waste showed higher diversity in the bacterial community.

To compare the relationship between bacterial communities in all the samples at the genus level, PCoA (Principal Coordinate Analysis) and UPGMA (unweighted pair group method with arithmetic mean) clustering based on the Bray-Curtis dissimilarity index were performed. The beta diversity indices, based on PCoA, revealed clear distinctions between different water samples forming three distinctive clusters (Figure 1C). Cluster one was formed exclusively by natural stream sources and associated with wet taro field water samples irrespective of the sampling site except for sample R-F1-E. The second distinctive cluster was formed by water samples collected from North Oahu, S-S3-N, T-S6-N, and R-S7-N, except S-F3-N. Interestingly, the 3rd cluster was formed by field water samples R-F1-E and S-F3-N indicating a close association between their bacterial communities, despite having been surveyed from different geographical locations and irrigated by different water sources (spring and river sources). Furthermore, UPGMA clustering revealed a similar clustering pattern in the dissimilarity of relative abundance of the bacterial communities (Supplementary Fig. 3). To unravel the close microbial association between R-F1-E and S-F3-N, these two samples were further sequenced to obtain full length 16S RNA and metagenomes using Oxford Nanopore MinION and Illumina NovaSeq, respectively, for amplicon and functional analyses.

### Full length 16S RNA amplicon analysis—Oxford Nanopore MinION

Samples R-F1-E and S-F3-N were sequenced in duplicate to attain confidence and reliability in the obtained data (Supplemental Table 6). Replicate 1 of sample S-F3-N failed to sequence and no reads were generated; nevertheless, the other replicate generated 87,818 reads with ∼1,500 bp length. In contrast, sample R-F1-E sequenced in two repeats validly sequenced 1,27,647 and 5,57,290 reads ranging from 1,500 to 1,600 bp length, and the comparative analyses between replicates at the genus and species levels were comparable, comprising almost similar bacterial composition (Supplementary Figure 4). Therefore, for further comparative analysis, reads from one sequencing replicate of sample R-F1-E were used.

### Taxonomic classification at phylum, genus, and species levels

At the phylum level, sample R-F1-E showed Bacteroidetes and Proteobacteria with relative abundance of >1%, while sample S-F1-E was dominated with 3 phyla-Bacteroidetes, Proteobacteria and Verrucomicrobia (Figure 2). Classification at the genus level uncovered a total of 11 and 6 genera from samples R-F1-E and S-F3-N, respectively, with relative abundance >1% (Figure 2). The most abundant genus classified in both the samples was *Limnohabitans* belonging to the family Comamonadaceae. Within the family Comamonadaceae, the genera *Arcobacter*, *Curvibacter*, *Limnohabitans*, and *Rhodoferax* were identified in both samples, with an additional two genera—*Hydrogenophaga* and *Pelomonas—*exclusively in sample R-F1-E with >1% relative abundance. Furthermore, genus *Aquirufa* was recognized in sample S-F3-N with relative abundance of 25.71%, while 8.86% remained in sample R-F1-E. The bacterial genera classified with relative abundance of <1% in total comprised 33.21% and 22.41% of bacterial community in sample R-F1-E and S-F3-N, respectively.

**Figure 2.**
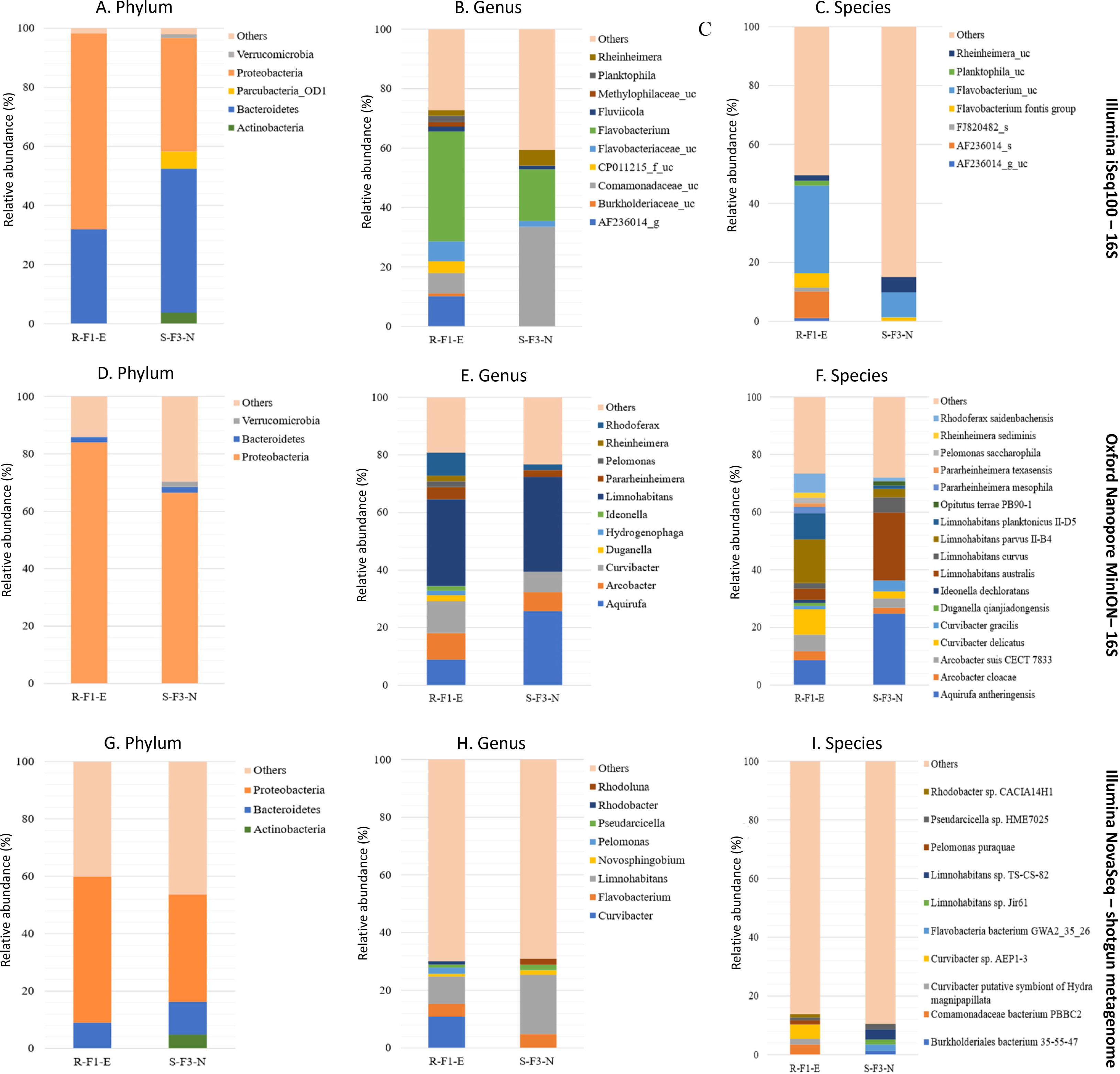
Comparison of sample R-F1-E and S-F3-N sequenced using Illumina iSeq100 (short amplicon reads), Oxford Nanopore MinION (long amplicon reads), and Illumina NovaSeq (shotgun reads) for the classification of phylum (**A**, **D**, and **G**, respectively), genus (**B**, **E**, and **H**, respectively), and species (**C**, **F**, and **I**, respectively) with relative abundance >1%. “Others” in the plots represents reads classified with <1% relative abundance and reads that remains unclassified.

At the species level, 16 and 11 species were classified from samples R-F1-E and S-F3-N, respectively, with >1% relative abundance (Figure 2F). Samples R-F1-E and S-F3-N were dominated with species *Limnohabitans parvus* II-B4 and *Aquirufa anthreingensis,* respectively. Four species belonging to genus *Limnohabitans*—*L. australis*, *L. curvus*, *L. parvus* II-B4, and *L. planktonicus—*were identified in both the samples with variable abundance. Furthermore, 73.38% and 72.06% of the bacterial diversity was composed of the bacterial population identified with relative abundance >1% in samples R-F1-E and S-F3-N, respectively. Full length amplicon reads that remained unclassified in samples R-F1-E and S-F3-N were 1.1 and 0.82% of the total analyzed reads, respectively.

### Taxonomic classification comparison with short and long reads 16S rRNA-based data sets

Short and full length 16S rRNA amplicon reads were obtained using Illumina iSeq100 and Oxford Nanopore MinION sequencers. The taxonomic classification results at phylum, genus and species levels were compared with different input reads (10K, 20K, 30K, 40K, and 50K), randomly extracted from total obtained valid reads–for samples R-F1-E and S-F3-N (Figure 3).

**Figure 3.**
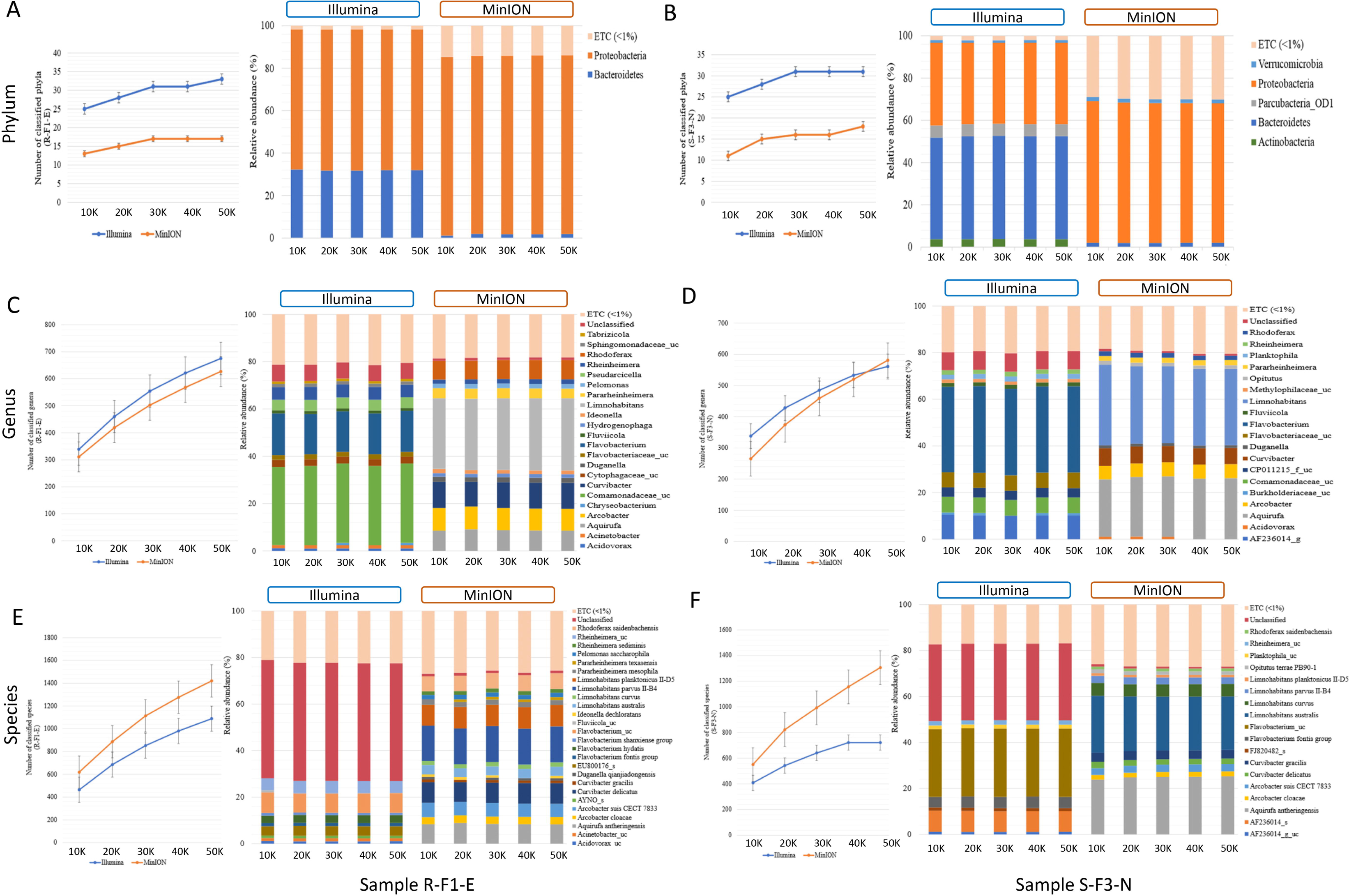
Comparison of **A)** total number of classified phyla; **B)** the phyla classified with >1% relative abundance; **C)** total number of classified genera; **D)** genus classified with >1% relative abundance; **E)** total number of classified species; and **F)** species classified with >1% relative abundance from sample R-F1-E and S-F1-E sequenced using Illumina iSeq100 and Oxford Nanopore MinION at different input reads ranging from 10K to 50K. “ETC (<1%) represents the classified reads at different taxonomic levels with <1% relative abundance, whereas “unclassified” represents the relative abundance of the reads that remains unclassified at taxonomic level.

At phylum level classification, Illumina sequenced samples R-F1-E and S-F3-N identified a greater number of phyla than MinION at different input reads (Figure 3A). In sample R-F1-E, an increase in the number of identified phyla was observed from 10K to 20K reads sequenced using Illumina (25 and 28, respectively) and MinION (13 and 15, respectively). With an increase in Illumina and MinION reads from 30K to 50K, a uniform number of phyla were identified, except for Illumina sequenced input read of 50K (Figure 3A). A similar trend in the number of identified phyla was observed in sample S-F3-N, with an exception that uniformity in the number of identified phyla (31) was observed in Illumina sequenced reads from 30K to 50K (Figure 3B). However, MinION sequenced input reads of 30K to 40K identified 16 phyla with a slight increase to 18 at 50K reads. Proteobacteria and Bacteroidetes were two major phyla identified in sample R-F1-E with >1% relative abundance, sequenced using both the techniques (Figure 3A). However, in sample S-F3-N, total 5- Actinobacteria, Bacteroidetes, Parcubacteria_OD1, Proteobacteria and, Verrucomicrobia and 3-Bacteroidetes, Proteobacteria and Verrucomicrobia were identified with relative abundance >1% from Illumina and MinION sequenced reads, respectively, at different input reads (Figure 3B). The number of genera and the genera classified with relative abundance >1% and remaining unclassified reads formed a uniform trend using both short- and long-amplicons at different input reads. The number of genera identified using Illumina input reads from 10K to 50K ranged from 339 to 675 for sample R-F1-E, whereas ranged from 338 to 561 for sample S-F3-N (Figure 3C and 3D). In contrast, MinION sequenced reads identified comparatively fewer genera ranging from 311 to 627 and 265 to 581 for sample R-F1-E and S-F3-N, respectively (Figure 3C and 3D). However, most genera classified using short amplicon reads were identified with low confidence values against the database, whereas long amplicon reads had comparatively better resolution for classified genera (Figure 3C and 3D). For both samples, the unclassified reads were fewer than 8% and 2% of the total input reads using short and long amplicon reads, respectively.

The number of species classified using long amplicon reads was higher than when using short amplicon reads (Figure 3). The number of identified species ranged from 619 to 1,421 and 551 to 1,306 for MinION sequenced samples R-F1-E and S-F3-N, respectively (Figure 3E and 3F). Whereas Illumina sequenced samples R-F1-E and S-F3-N identified species ranging from 464 to 1089 and from 408 to 722, respectively (Figure 3E and 3F). At the species level classification, ∼50% and ∼33% of the total input reads remained unclassified using short amplicon reads for sample R-F1-E and S-F3-N, respectively, whereas long amplicon reads were classified with high accuracy comprising >98% classified reads (Figure 3E and 3F). In sample R-F1-E and S-F3-N, the species identified with relative abundance >1%, utilizing long amplicon reads at different inputs comprehends >70% of the identified bacterial microbiota.

In term of relative abundance, almost similar abundance patterns were obtained with each technique when 10 – 50K reads were used as an input data—indicated that minimum input of 10K reads from either Illumina iSeq100 or Oxford Nanopore MinION, can provide similar resolution with 5 times more input reads. However, with respect to the number of classified phyla, Illumina provided better outcomes compared to Oxford Nanopore, and there was no dramatic increase in number of phyla when the input reads were increased from 10K to 50K by either sequencing technology (Figure 3A and 3B). The analyses indicated that Oxford Nanopore MinION is a better choice for higher resolution at genus and species levels (Figure 3C-3F). To identify number of genera or species, it is important to include higher number of reads (∼>20K).

### Shotgun metagenome analysis

A total of 5,61,183 and 4,91,726 non-redundant genes were identified from sample R-F1-E and S-F3-N, respectively, while sharing 1,24,661 (12%) unigenes between both. Despite having close microbial association indicated by PCoA analysis, the samples R-F1-E and S-F3-N were distinctively differentiated based on unique genes composition of 78% and 75%, respectively.

### Taxonomic classification of metagenomics (shotgun) data

According to the obtained abundance table of each taxonomic level, the bar plots were plotted for the top 10 classified phyla, genera, and species (Figure 2G-2I). At the phylum level, the most abundant phyla, in both the samples, were Proteobacteria, followed by Bacteroidetes with relative abundance >1%. Additionally, Actinobacteria was also classified in sample S-F3-N with >1% relative abundance, differentiating this from sample R-F1-E in which seven genera—*Curvibacter, Limnohabitans, Flavobacterium, Pelomonas, Rhodobacter, Pseudarcicella,* and *Novosphingobium—*were classified with more than 1% relative abundance, whereas only 5 genera—*Limnohabitans, Flavobacterium, Rhodoluna, Pesudarcicella,* and *Novosphingobium—*were classified in sample S-F3-N. Species level classification revealed 10 species with relative abundance of >1% from both the samples. A high percentage of “others” in the metagenomic analysis could result from an incomplete database.

“Others” representing the relative abundance of the reads that remain unclassified and classified with relative abundance of <1% was higher at phylum, genus, and species level classification for both the samples sequenced using Illumina NovaSeq (shotgun reads) than Illumina iSeq100 and Oxford Nanopore MinION (Figure 2). Sample R-F1-E represented 40.12%, 69.88%, and 86.12% of reads as “others” at phylum, genus, and species level classification, respectively. Sample S-F3-N at phylum, genus, and species level represented 46.25%, 69.04%, and 89.53%, respectively, as “others”.

### Functional profiling of active bacterial community

For better insight into the physiology of a bacterial community, the assembled metagenomic protein coding sequences were mapped against three functional databases—eggNOG, KEGG, and CAZy (Supplemental Figure 4). Both samples (R-F1-E and S-F3-N) revealed similarity in annotated gene function profiles and were clustered together.

Annotation based on eggNOG database revealed (Supplementary Fig. 5A-B) that highest number genes in sample R-F1-E were associated with inorganic ion, amino acid, carbohydrate, nucleotide, and lipid transport and metabolism, cell motility, and transcription with the relative abundance >1% for each function. Whereas in sample S-F3-N, the maximum number of genes were associated with 7 functions and having >1% relative abundance—replication, recombination, and repair, translation, ribosomal structure, and biogenesis, nucleotide transport and metabolism, cell wall/membrane/envelope biogenesis, post-translational modification, protein turnover, chaperons, coenzyme transport and metabolism, and energy production and conversion.

Most of the genes represented in the KEGG pathway analysis were associated with metabolic pathways (Supplemental Fig. 4C-D), and particularly dominant in the category of amino acid transport and metabolism having 28,924 and 19,900 associated genes in samples R-F1-E and S-F3-N, respectively. Statistically differential features of functional categories based on KEGG analysis between the two samples were analyzed using STAMP, indicating metabolism, genetic information processing, human diseases, and organismal system dominant in sample S-F3-N, whereas environmental information and cellular processing were enriched in sample R-F1-E (Supplemental Fig. 4D).

**Figure 4.**
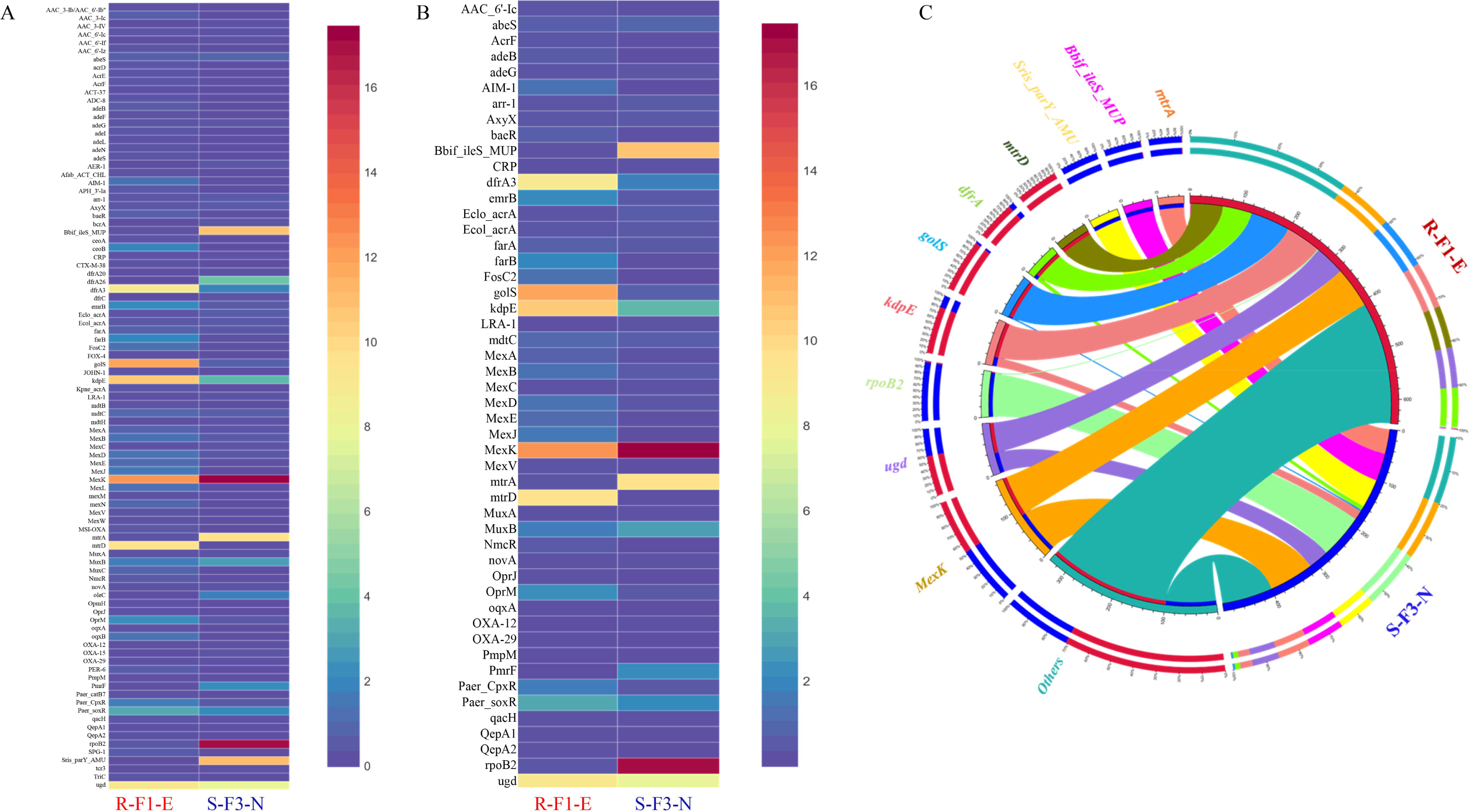
Distribution heatmap to represent **A)** comparison of relative abundance of a total 95 Antibiotic resistance gene (ARG) profile obtained from sample R-F1-E and S-F3-N; **B)** comparison of relative abundance of 50 ARGs shared between sample R-F1-E and S-F3-N. All the unique genes from the metagenomic assembly were blastp against Comprehensive Antibiotic Resistance Database (CARD). **C)** Circos analysis displays the corresponding abundance relationship between samples and top 10 identified antibiotic resistance genes (ARGs) along with “others” representing remaining ARGs. Circle chart is divided into two parts. The right side of the circle is sample information, and the left side of the circle represents top 10 ARGs. Inner circle with different colors represents different ARGs. The scale represents the relative abundance, and the unit is ppm. The left part represents the sum of relative abundance of different samples for ARG, while the outer right circle represents the relative abundance of different ARGs in the samples.

As per CAZy database-based analysis, glycoside hydrolases (GH) associated genes were most abundant with the relative abundance of 49.33 and 51.87% in sample R-F1-E and S-F3-N, respectively, followed by glycosyl transferase (GT), carbohydrate-binding modules (CBM), carbohydrate esterases (CE), auxiliary activities (AA), polysaccharide lyases (PL) (Supplementary Fig. 5E). STAMP analysis revealed GH was significantly different with a q-value of 4.37e-3 and was enriched in sample S-F3-N (Supplementary Fig. 5F). Whereas glycosyl transferase (GT), carbohydrate-binding modules (CBM), carbohydrate esterases (CE), auxiliary activities (AA), polysaccharide lyases (PL) were higher in sample R-F1-E, with no significant differences observed among these functions.

### Occurrence, abundance, and diversity of ARGs

To explore and compare the ARGs profile in sample R-F1-E and S-F3-N, all unique genes obtained from the samples were BLASTp against the CARD database. This analysis revealed the presence of 83 and 62 ARGs in sample R-F1-E and S-F3-N, respectively (Figure 4A), while sharing 50 ARGs between each other with variable relative abundance (Figure 4B). *MexK,* a resistance nodulation cell division (RND) antibiotic efflux pump gene, was the most abundant ARG present in both the samples (Figure 4C).

Furthermore, the top 10 most abundant ARGs out of 95 ARGs, annotated collectively from both samples, were represented in Circos for observing overall proportion and distribution of the resistance genes in both samples (Figure 4C). The top 10 ARGs were: *mexK* (multidrug resistance gene), *ugd* (peptide resistance gene), *rpoB2* (rifamycin resistance gene), *kdpE* (aminoglycoside resistance gene), *golS* (multidrug resistance gene), *dfrA3* (diaminopyrimidine resistance gene), *mtrD* (macrolide resistance gene), *Streptomyces rishiriensis parY* mutant conferring resistance to aminocoumarin (*Sris_parY_AMU*) (aminocoumarin resistance gene), Bifidobacterium *ileS* conferring resistance to mupirocin (*Bbif_ileS_MUP*) (mupirocin resistance gene), and *mtrA* (macrolide resistance gene). The relative abundance of gene *ugd*, *kdpE*, *golS*, and *dfrA3* was higher in sample R-F1-E, whereas *mexK*, *rpoB2*, *Bbif_ileS_MUP*, and *mtrA* were relatively higher in sample S-F3-N. Interestingly, ARG *mtrD* and *Sris_parY_AMU* were only conferred to sample R-F1-E and S-F3-N, respectively.

An additional analysis was performed to reveal the dominant bacterial phyla possessing the most ARG genes with different associated resistance mechanisms. The most abundant resistant mechanism associated with the annotated ARGs corresponded to RND antibiotic efflux pump, followed by major facilitator superfamily (MFS) antibiotic efflux pump, antibiotic target alteration (pmr phosphoethanolamine transferase), protein and two component regulatory system modulating antibiotic efflux (*kdpE*), antibiotic target replacement (*DfrA42_TMP*), and ABC antibiotic efflux pump. These potential antibiotic mechanisms were associated with the ARG that were affiliated with phyla Proteobacteria (Supplementary Fig. 6).

## DISCUSSION

Our study highlighted significant differences and similarities in the bacterial communities of different irrigation water systems from different geographical locations (North, West, and East) on Oahu, Hawaii. Comparative assessment of bacterial communities between samples showed distinctive discriminations based on type of water system and geographical location. It is striking to note that natural stream and associated field water samples were dominated by Proteobacteria, regardless of their geographical locations—there was a close bacterial association between the samples based on beta diversity analysis. These outcomes agreed with the previous studies conducted in Brazil (36) and Tokyo (37), which revealed a dominance of Proteobacteria in river water. Samples collected from North Oahu showed close microbial association regardless of different water systems, indicating an influence of geographical locations (topography, water bodies, climatic conditions, natural vegetation etc.) in composing the microbial consortia (38).

Field water samples R-F1-E and S-F3-N were clustered based on the microbiota despite being irrigated by different irrigation systems (spring and stream) and different geographical regions (North and East), which prompted us to uncover the complex and diverse microbiota at a higher taxonomic level (Figure 1). The short amplicon reads generated from V3-V4 gene region of 16S rRNA using Illumina iSeq100 was able to detect phyla with high accuracy in addition to classification of most dominant genera as well. However, some genera within the family were not classified with high confidence value and more than 50% of the valid reads were unclassified, indicating a limitation of short amplicon reads for high resolution and accuracy of classification. A study (39) designed to uncover and compare the microbial consortia of indoor dust sequenced using Illumina and Nanopore MinION revealed significant differences in microbial composition at genus and species levels, with better resolution provided by MinION sequenced reads. Therefore, to investigate the microbiota of sample R-F1-E and S-F3-N at a higher taxonomic level with better resolution, full length 16S rRNA gene region was sequenced using Oxford Nanopore MinION and analyzed. Full length amplicon analysis revealed high abundance of the genus *Limnohabitans* that includes planktonic bacteria and classified other dominant genera within family Comamonadaceae that remained unclassified using short amplicon reads. All the four species within the genus *Limnohabitans* (40, 41) were successfully classified with >1% relative abundance. Additionally, genus *Aquirufa*, a freshwater bacterium, was identified in spring and stream field water with relative abundance >1% and *Aquirufa antheringensis* was the dominant species in spring field water. Another study (42) also found the higher abundance of *A. antheringensis* in fresh water. The resolution obtained for genus and species level classification was better using long amplicon reads with <2% valid reads that remained unclassified (Figure 2).

Furthermore, we compared the performance of long reads (∼1,500bp) obtained from Oxford Nanopore MinION with short reads (∼300bp) obtained from Illumina iSeq100 to assess bacterial taxonomic classification at phylum, genus, and species levels with different numbers of input reads. Results from this experimental study showed uniform trends in classification at phylum, genus, and species levels for samples, R-F1-E and S-F3-N, at 10K, 20K, 30K, 40K, and 50K input reads (Figure 3). However, when long-and short-read outcomes were compared, dissimilarities in relative abundance at all three taxonomic levels were observed (Figure 3).

Short-read-based taxonomic analysis provided the most comprehensive classification at the phylum level compared to 16S rRNA full length reads and shotgun metagenome data (Figures 2-3). However, 16S rRNA full length reads clearly illustrated its advantage for classification at genus and species levels (Figures 2-3). In a study (43) proposed Oxford Nanopore MinION as a low cost and rapid technology for revealing microbial communities with higher resolution at the species level which ultimately aids in identifying bacteria potentially pathogenic to human health. In our study, with a high number of unclassified reads at phylum [39.52% (R-F1-E); 45.82% (S-F3-N)], genus [68.04% (R-F1-E); 68.35% (S-F3-N)] and, species [85.37% (R-F1-E);

89.17% (S-F3-N)] levels, we have not observed any advantages of using shotgun metagenome data for taxonomic classification (Figure 2)—this could be due to the limited and incomplete annotated metagenomic and bacterial genome databases currently available (44). With the advancement and improvement in the Nanopore MinION technology, this efficient, cost-effective, and robust technology can be employed for on-field microbiome study of environmental samples with minimum data requirements (45).

The environmental samples consist of complex and diverse microbiota which are better resolved in terms of predication of microbial community’s functions. This can be achieved using shotgun metagenomic sequencing with advanced next generation sequencing technologies that generates enormous amounts of genomic data (46). However, due to different sequencing protocols and annotated databases, metagenome analysis and 16S rRNA gene sequencing cannot provide an identical taxonomic classification, as observed in our study and in (47). Metagenomic functional analysis revealed the presence of 78% and 75% of unique genes in sample R-F1-E and S-F3-N, respectively, while only 12% of the genes were shared between both the samples, but interestingly, were annotated for comparable gene functional profiles (Supplementary Fig. 4).

The relatively high abundance of genes was related to metabolism of amino acids, nucleotides, carbohydrates, coenzymes, lipids, and inorganic ion metabolism and transport. ‘Amino acid metabolism’ was enriched in both the samples, which may be due to fertilizer residues that provide a suitable living environment for microbiota that use amino acids. Additionally, environmental samples consist of diverse and abundant complex mixtures of carbohydrates requiring different enzymes for metabolism, mainly supported by glycoside hydrolases (GH) (48). In our study, GH were the most abundant and significantly different among all the other identified enzymes in both samples (Figure 4E and 4F). This enzyme assists in the enzymatic processing of carbohydrate, ultimately contributing to functioning of an ecosystem, global carbon cycling. The metagenomic data also revealed the prevalence of a variety of ARGs in both the samples. The ubiquity of ARGs in the environmental sample is an emerging concern. A study (49) documented the prevalence of ARGs in irrigation ditch water and urban/agriculturally impacted river sediments leading to the potential spread of ARGs to or from humans. From 95 identified ARGs, only 50 genes were shared between both the samples with variable abundance depending on the microbial consortia and their genome compositions (Figure 4)—the genomic composition can be altered through horizontal gene transfer from environment or other bacteria mediated by mobile genetic elements such as plasmids, transposons, bacteriophages, insertion sequences and integrons (50, 51). The most abundant ARG in both the samples was *MexK*, a resistance nodulation cell division (RND) antibiotic efflux pump gene which can transport multiple classes of antimicrobials, contributing to multidrug resistance (52). Therefore, uncovering the bacterial components, functional analysis, and investigation of the ARGs will resolve the microbial complexity and help to formulate better disease management strategies for water transmitted pathogens.

## CONCLUSIONS

The bacterial consortia found in different water source of taro irrigation across the island of Oahu, Hawaii revealed that Proteobacteria is the most dominant phyla, except for a few samples from storage tank and spring water. The most reliable and comprehensive taxonomic classifications at phylum and genus/species levels were observed with input reads obtained from Illumina and Oxford Nanopore, respectively. The lack of robust and comprehensive annotated metagenome and bacterial genome databases contributed to inconclusive classification using shotgun metagenome reads, particularly at genus and species levels. However, metagenomic data contributed to the understanding of gene distribution of microbiomes and their functions, including ARGs, associated with different microbial consortia. This study provided some appropriate sequencing platforms and pipelines to study irrigation water microbiome.

## ACKNOWLEDGEMENTS

This research was supported in parts by NIGMS of the National Institutes of Health (P20GM125508), USDA National Institute of Food and Agriculture, and College of Tropical Agriculture and Human Resources managed Hatch project (9038H).

## SUPPLEMENTARY FIGURES AND TABLES

**Supplementary Figure 1.** Bar plot comparison of phylum level classification, classified with relative abundance of >1% in 11 samples-R-F1-E, R-S1-E, R-S2-W, R-F2-W, S-F3-N, R-S4-SE, R-F4-SE, R-S5-SE, R-F5-E, T-S6-N, and R-S7-N (Replicate 1 and Replicate 2) sequenced for short length amplicon using Illumina iSeq100 and analyzed on EzBioCloud platform. “Others” represents the reads classified with less than <1% relative abundance and remains unclassified in the classification against the database.

**Supplementary Figure 2.** Distribution heatmap of bacterial species classified with >1% relative abundance among all the 12 water samples—sequenced for V3-V4 region of 16S rRNA gene region using Illumina iSeq100 sequencing platform. The generated short amplicon reads were analyzed using EzBioCloud platform. The heatmap was generated using displayR.

**Supplementary Figure 3.** UPGMA (unweighted pair group method with arithmetic mean) clustering of water samples based on Bray-Curtis dissimilarity index at genus level. Samples were grouped in three distinctive clusters: Cluster 1 (R-F1-E and S-F3-N) irrespective of water system or geographical location, Cluster 2 (R-S1-E, R-F2-W, R-S2-W, R-F4-SE, R-S4-SE, R-F5-SE, and R-S5-SE) based on irrigation source and associated taro field water, and Cluster 3 (S-S3-N, T-S6-N, and R-S7-N) based on geographical location.

**Supplementary Figure 4.** Bar plot comparing the (A) genus and (B) species classified with relative abundance of >1% in sample R-F1-E (Replicate 1 and Replicate 2) sequenced for full length amplicon using Oxford Nanopore MinION and analyzed on EPI2ME platform. Input valid reads that were not classified to genus and species levels are represented as “Unclassified”, while “ETC (<1%)” represents the bacterial population identified with relative abundance of <1%.

**Supplementary Figure 5.** Comparison of samples R-F1-E and S-F3-N for relative abundance and statistical differences of annotated gene function profiles based on mapping of assembled metagenomic protein coding sequences to three databases: (A, B) non-supervised Orthologous groups (eggNOG), (C, D) Kyoto Encyclopedia of Genes and Genomes (KEGG), and (E, F) Carbohydrate-Active Enzymes Database (CAZy). Statistical analyses performed using STAMP v2.1.3 software, employing Fisher’s exact test with Newcombe-Wilson CI method and Benjamini-Hochberg FDR correction factors, and visualized using extended error bar plots.

**Supplementary Figure 6.** Circos analysis displays the corresponding abundance relationship between identified dominant phyla (Proteobacteria and Actinobacteria) along with “other” representation of identified phyla and associated resistance mechanism. Circle chart is divided into two parts. The right side of the circle is phyla information, and the left side of the circle is antibiotic resistance mechanisms. Inner circle with different colors represents different antibiotic resistance mechanisms. The scale represents the relative abundance, and the unit is ppm. The left part represents the sum of relative abundance of different phyla for resistance mechanisms, while the outer right circle vice versa.

**Supplemental Table 1.** List of samples sequenced in two replicates using Illumina iSeq100. Assigned barcodes with different combinations of i5 and i7 adapters.

**Supplemental Table 2.** List of valid reads with calculated average read length generated by sequencing of each barcode after quality filtration.

**Supplemental Table 3** Statistical analysis of the identified phyla among all the samples was determined using one-way ANOVA (single factor) with the least significant difference (LSD) test at α=0.05.

**Supplemental Table 4.** Statistical analysis of identified genera among all the samples was determined using one-way ANOVA (single factor) with the least significant difference (LSD) test at α=0.05.

**Supplementary Table 5.** Short length 16S rRNA reads classified to genus level, accounting for relative abundance <1% and remains unclassified are represented as “ETC (<1%)” and “Unclassified” based on the analysis performed using EzBioCloud.

**Supplemental Table 6.** Oxford Nanopore MinION 16S rRNA sequencing and analyses results of sample R-F1-E and S-F3-N. The EPI2ME Fastq16S pipeline was used for the analyses.

