## Supplementary material for "Exploring Taxonomic and Functional Microbiome of Hawaiian Stream and Spring Irrigation Water Systems Using Illumina and Oxford Nanopore Sequencing Platforms": Supplemental Materials.pdf

### SUPPLEMENTARY FIGURES AND TABLES

**Supplementary Figure 1.** Bar plot comparison of phylum level classification, classified with relative abundance of >1% in 11 samples- R-F1-E, R-S1-E, R-S2-W, R-F2-W, S-F3-N, R-S4-SE, R-F4-SE, R-S5-SE, R-F5-E, T-S6-N, and R-S7-N (Replicate 1 and Replicate 2) sequenced for short length amplicon using Illumina iSeq100 and analyzed on EzBioCloud platform. "Others" represents the reads classified with less than <1% relative abundance and remains unclassified in the classification against the database.

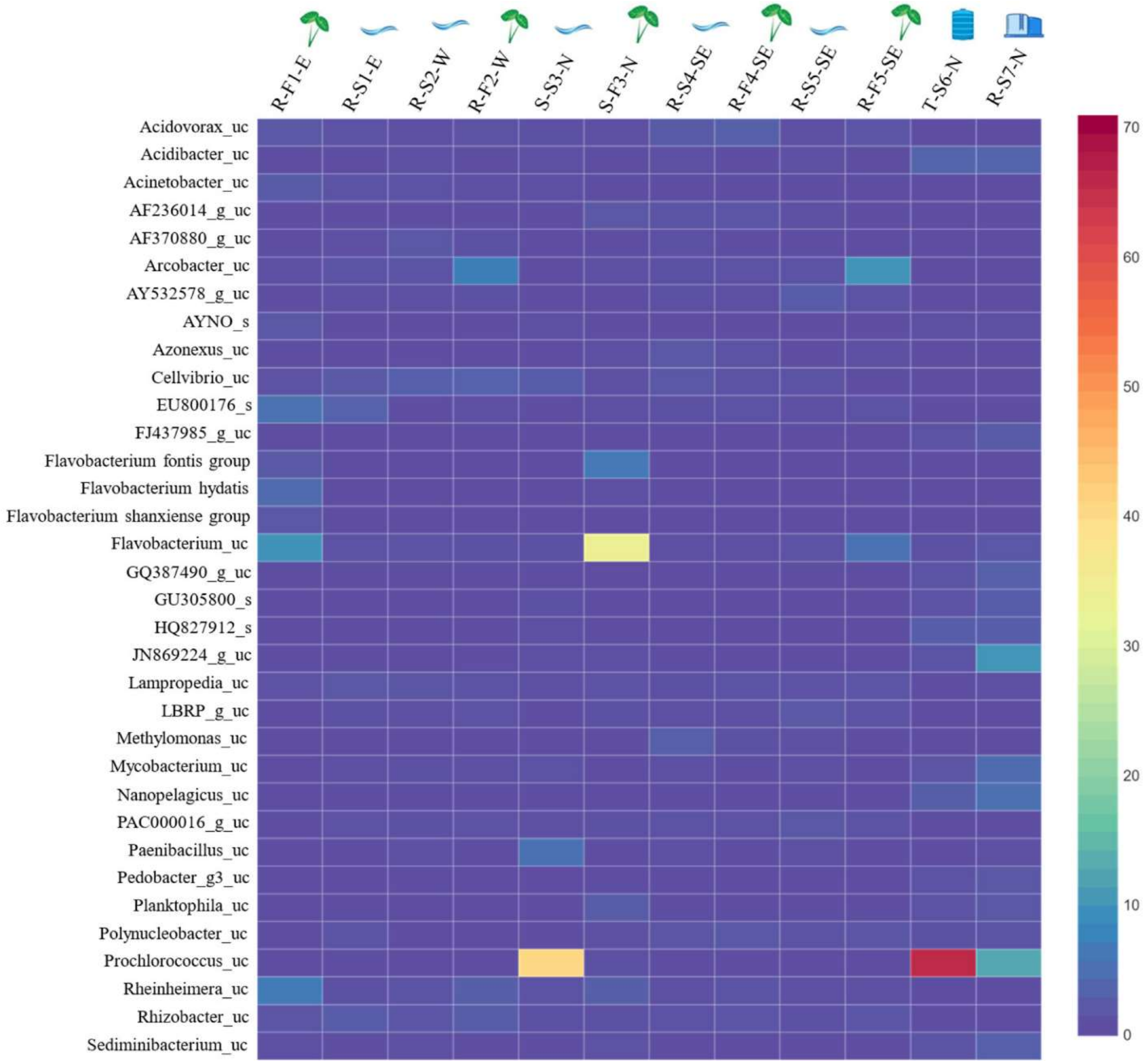

**Supplementary Figure 2.** Distribution heatmap of bacterial species classified with >1% relative abundance among all the 12 water samples—sequenced for V3-V4 region of 16S rRNA gene region using Illumina iSeq100 sequencing platform. The generated short amplicon reads were analyzed using EzBioCloud platform. The heatmap was generated using displayR.

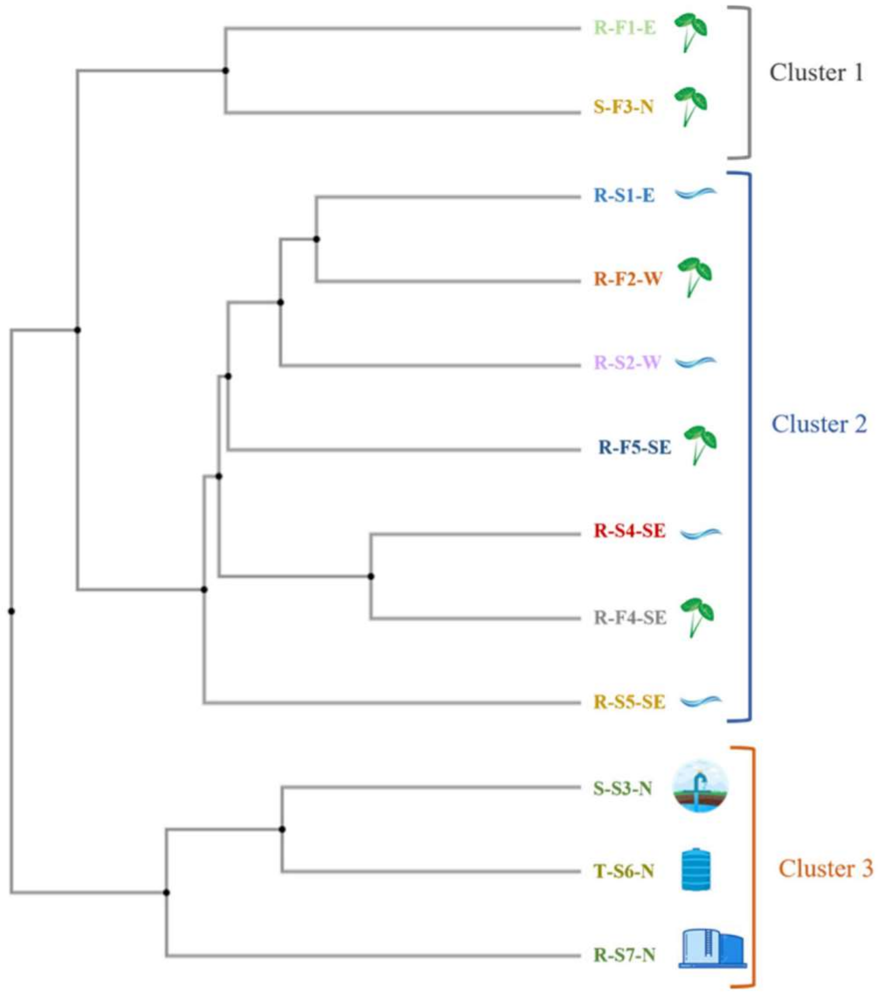

**Supplementary Figure 3**. UPGMA (unweighted pair group method with arithmetic mean) clustering of water samples based on Bray-Curtis dissimilarity index at genus level. Samples were grouped in three distinctive clusters: Cluster 1 (R-F1-E and S-F3-N) irrespective of water system or geographical location, Cluster 2 (R-S1-E, R-F2-W, R-S2-W, R-F4-SE, R-S4-SE, R-F5-SE, and R-S5-SE) based on irrigation source and associated taro field water, and Cluster 3 (S-S3-N, T-S6-N, and R-S7-N) based on geographical location.

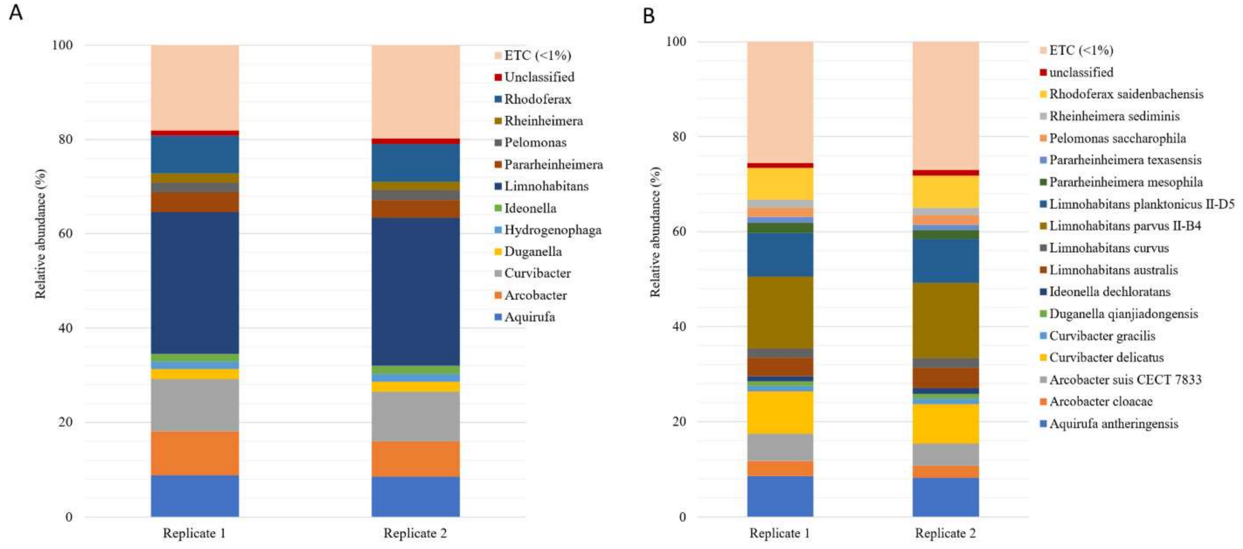

**Supplementary Figure 4**. Bar plot comparing the (A) genus and (B) species classified with relative abundance of >1% in sample R-F1-E (Replicate 1 and Replicate 2) sequenced for full length amplicon using Oxford Nanopore MinION and analyzed on EPI2ME platform. Input valid reads that were not classified to genus and species levels are represented as “Unclassified”, while “ETC (<1%)” represents the bacterial population identified with relative abundance of <1%.

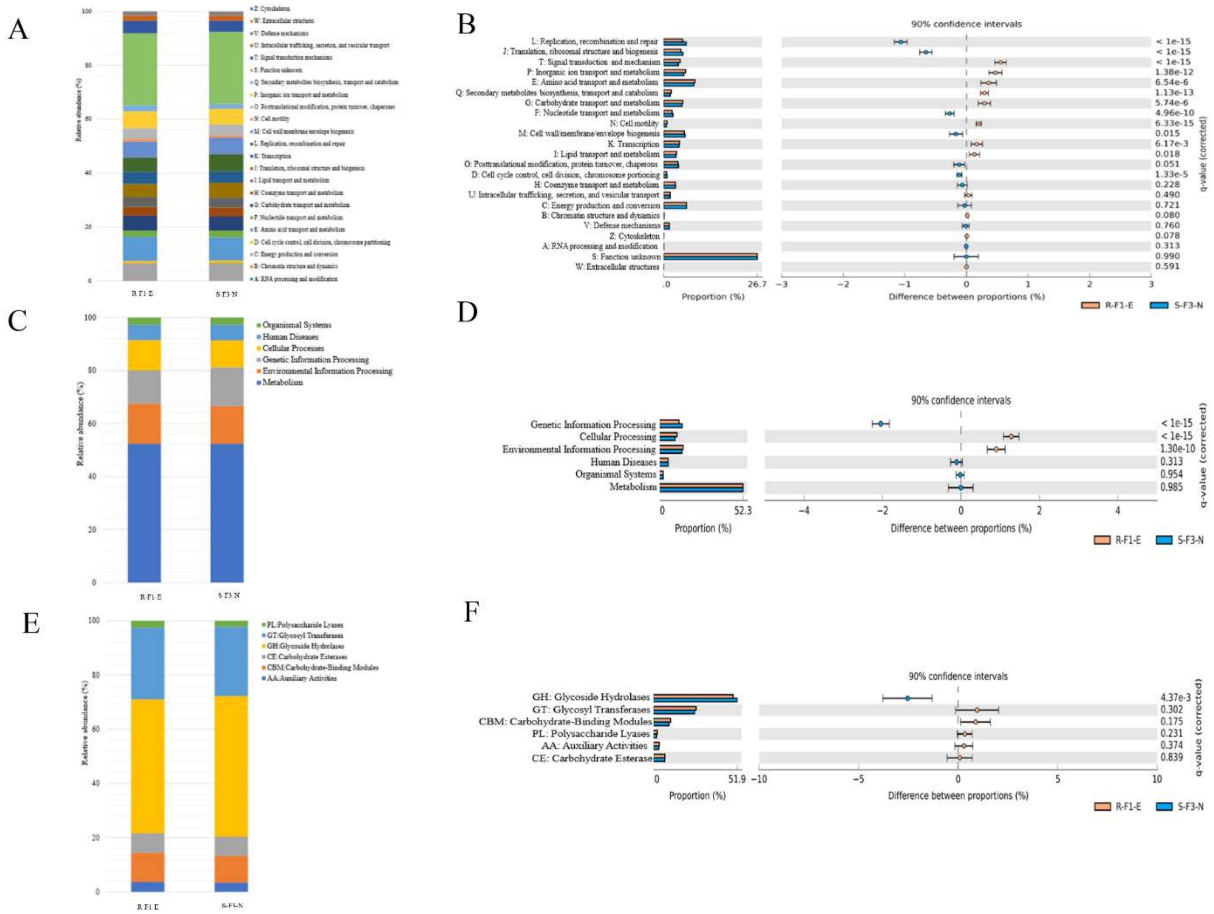

**Supplementary Figure 5.** Comparison of samples R-F1-E and S-F3-N for relative abundance and statistical differences of annotated gene function profiles based on mapping of assembled metagenomic protein coding sequences to three databases: (A, B) non-supervised Orthologous groups (eggNOG), (C, D) Kyoto Encyclopedia of Genes and Genomes (KEGG), and (E, F) Carbohydrate-Active Enzymes Database (CAZy). Statistical analyses performed using STAMP v 2.1.3 software, employing Fisher's exact test with Newcombe-Wilson CI method and Benjamini-Hochberg FDR correction factors, and visualized using extended error bar plots.

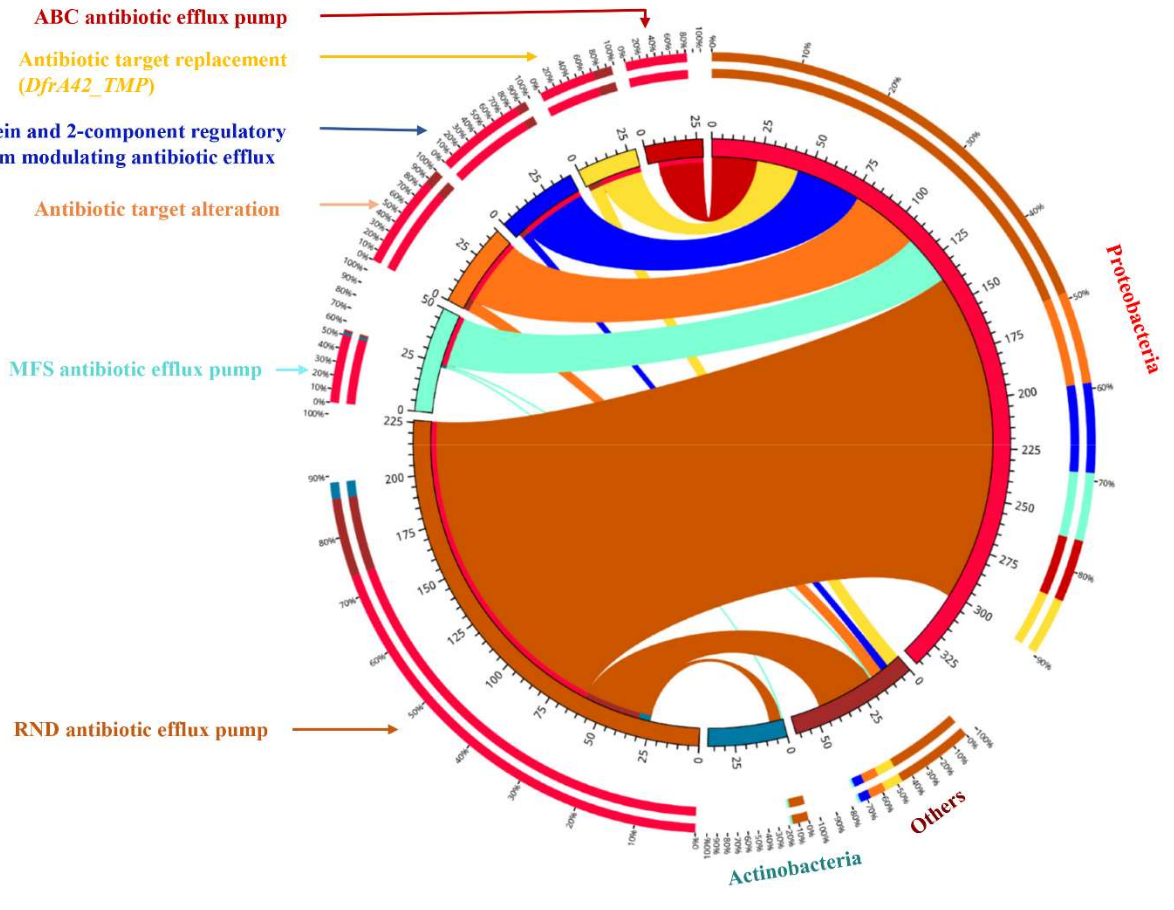

**Supplementary Figure 6**. Circos analysis displays the corresponding abundance relationship between identified dominant phyla (Proteobacteria and Actinobacteria) along with "other" representation of identified phyla and associated resistance mechanism. Circle chart is divided into two parts. The right side of the circle is phyla information, and the left side of the circle is antibiotic resistance mechanisms. Inner circle with different colors represents different antibiotic resistance mechanisms. The scale represents the relative abundance, and the unit is ppm. The left part represents the sum of relative abundance of different phyla for resistance mechanisms, while the outer right circle vice versa.

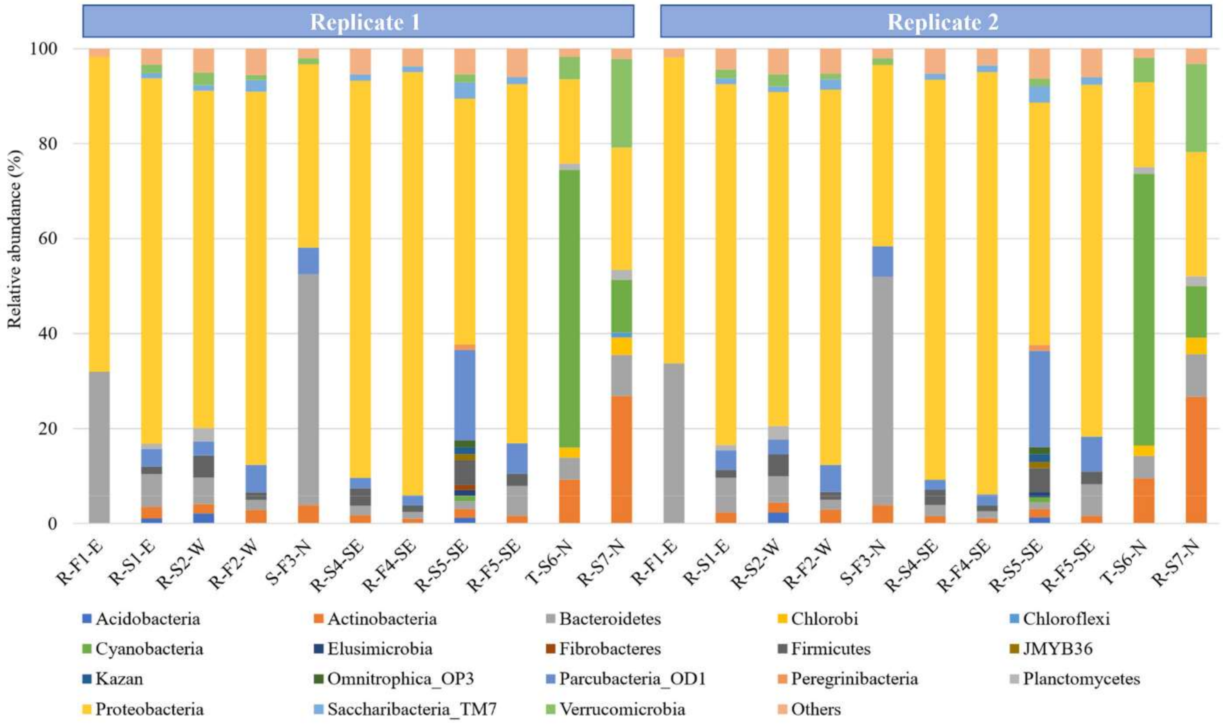

**Supplementary Table 1.** List of samples sequenced using Illumina iSeq100. Assigned barcodes with different combinations of i5 and i7 adapters. Each sample was sequenced in two replicates.

| Sample | Replicate 1 | Replicate 2 |
|---|---|---|
| R-F1-E | Barcode 1 (i517-i710 adapter) | Barcode 13 (i506-i710 adapter) |
| R-S1-E | Barcode 2 (i517-i705 adapter) | Barcode 14 (i506-i705 adapter) |
| R-S2-W | Barcode 3 (i517-i706 adapter) | Barcode 15 (i506-i706 adapter) |
| R-F2-W | Barcode 4 (i517-i707 adapter) | Barcode 16 (i506-i707 adapter) |
| S-S3-N | Barcode 5 (i517-i711 adapter) | Barcode 17 (i506-i711 adapter) |
| S-F3-N | Barcode 6 (i517-i714 adapter) | Barcode 18 (i506-i714 adapter) |
| R-S4-SE | Barcode 7 (i505-i710 adapter) | Barcode 19 (i503-i710 adapter) |
| R-F4-SE | Barcode 8 (i505-i705 adapter) | Barcode 20 (i503-i705 adapter) |
| R-S5-SE | Barcode 9 (i505-i706 adapter) | Barcode 21 (i503-i706 adapter) |
| R-F5-SE | Barcode 10 (i505-i707 adapter) | Barcode 22 (i503-i707 adapter) |
| T-S6-N | Barcode 11 (i505-i711 adapter) | Barcode 23 (i503-i711 adapter) |
| R-S7-N | Barcode 12 (i505-i714 adapter) | Barcode 24 (i503-i714 adapter) |

**Supplemental Table 2.** Valid reads after quality filtration with calculated average read length generated using illumine iSeq100 sequencer.

| Sample | Barcode | Valid reads | Average length (bp) | Barcode | Valid reads | Average length (bp) |
|---|---|---|---|---|---|---|
| R-F1-E | 1 | 50,796 | 286.8 | 13 | 47,573 | 286.6 |
| R-S1-E | 2 | 47,032 | 282.9 | 14 | 48,790 | 282.9 |
| R-S2-W | 3 | 61,272 | 283.2 | 15 | 67,325 | 283.2 |
| R-F2-W | 4 | 28,288 | 283.6 | 16 | 31,661 | 283.5 |
| S-S3-N | 5 | 47,654 | 282.4 | 17 | - | - |
| S-F3-N | 6 | 74,525 | 285.9 | 18 | 63,230 | 285.8 |
| R-S4-SE | 7 | 31,574 | 284.0 | 19 | 28,530 | 284.1 |
| R-F4-SE | 8 | 22,274 | 284.3 | 20 | 26,908 | 284.3 |
| R-S5-SE | 9 | 43,658 | 282.0 | 21 | 46,548 | 282.1 |
| R-F5-SE | 10 | 35,595 | 284.1 | 22 | 44,228 | 284.1 |
| T-S6-N | 11 | 47,020 | 281.2 | 23 | 55,980 | 281.3 |
| R-S7-N | 12 | 33,495 | 282.6 | 24 | 33,187 | 282.7 |

**Supplementary Table 3** Statistical analysis of the identified phyla among all the samples was determined using one-way ANOVA (single factor) with the least significant difference (LSD) test at α=0.05.

| Phylum | Count | Sum | Average | Variance |
|---|---|---|---|---|
| Acidobacteria | 12 | 8.0138 | 0.667817 | 0.459307 |
| Actinobacteria | 12 | 55.2168 | 4.6014 | 54.25061 |
| Bacteroidetes | 12 | 122.506 | 10.20883 | 215.7226 |
| Chlorobi | 12 | 7.0533 | 0.587775 | 1.321251 |
| Chloroflexi | 12 | 4.8577 | 0.404808 | 0.120637 |
| Cyanobacteria | 12 | 108.4629 | 9.038575 | 348.6479 |
| Elusimicrobia | 12 | 3.3087 | 0.275725 | 0.114638 |
| Fibrobacteres | 12 | 3.3986 | 0.283217 | 0.087266 |
| Firmicutes | 12 | 44.4733 | 3.706108 | 39.62349 |
| JMYB36 | 12 | 1.733 | 0.144417 | 0.15651 |
| Kazan | 12 | 4.4133 | 0.367775 | 0.178326 |
| Omnitrophica_OP3 | 12 | 3.5871 | 0.298925 | 0.174675 |
| Parcubacteria_OD1 | 12 | 48.4779 | 4.039825 | 26.67713 |
| Peregrinibacteria | 12 | 4.3685 | 0.364042 | 0.116108 |
| Planctomycetes | 12 | 11.6281 | 0.969008 | 0.646273 |
| Proteobacteria* | 12 | 702.3075 | 58.52563 | 641.4002 |
| Saccharibacteria_TM7 | 12 | 13.0376 | 1.086467 | 1.051675 |
| Verrucomicrobia | 12 | 36.3249 | 3.027075 | 25.42415 |

One way-ANOVA

| *Source of Variation* | *SS* | *df* | *MS* | *F* | *P-value* | *F crit* |
|---|---|---|---|---|---|---|
| Between Groups | 37621.02 | 17 | 2213.001 | 29.37238 | 5.63E-45 | 1.674587 |
| Within Groups | 14917.9 | 198 | 75.34293 | | | |
| Total | 52538.92 | 215 | | | | |

Significant if difference in variance between two compared phyla was greater than calculated LSD value (6.988063).

*Statistically significant among all other phyla identified.

**Supplementary Table 4.** Statistical analysis of identified genera among all the samples was determined using one-way ANOVA (single factor) with the least significant difference (LSD) test at α=0.05.

| Genus | Samples | Sum | Average | Variance |
|---|---|---|---|---|
| AF236014_g | 12 | 26.4928 | 2.207733 | 13.93174 |
| *Acidibacter* | 12 | 6.1776 | 0.5148 | 0.976789 |
| *Acinetobacter* | 12 | 2.5366 | 0.211383 | 0.151579 |
| *Arcobacter* | 12 | 16.9884 | 1.4157 | 7.655345 |
| *Azonexus* | 12 | 1.8406 | 0.153383 | 0.104587 |
| Burkholderiaceae_uc | 12 | 2.7073 | 0.225608 | 0.119841 |
| Flavobacteriaceae_uc | 12 | 9.3295 | 0.777458 | 3.698323 |
| GU305779_g | 12 | 3.1472 | 0.262267 | 0.339311 |
| *Mycobacterium* | 12 | 6.2091 | 0.517425 | 1.301162 |
| PAC000128_g | 12 | 2.146 | 0.178833 | 0.0872 |
| *Prevotella* | 12 | 2.8556 | 0.237967 | 0.155507 |
| *Prochlorococcus* | 12 | 105.1676 | 8.763967 | 351.5253 |
| Rhodocyclaceae_uc | 12 | 2.0787 | 0.173225 | 0.114571 |
| *Roseomonas* | 12 | 2.3434 | 0.195283 | 0.136651 |
| *Sulfuricurvum* | 12 | 1.4322 | 0.11935 | 0.089258 |
| *Tabrizicola* | 12 | 1.5307 | 0.127558 | 0.093728 |
| FJ437985_g | 12 | 2.0832 | 0.1736 | 0.206132 |
| *Flavobacterium* | 12 | 62.903 | 5.241917 | 123.6531 |
| *Hyphomicrobium* | 12 | 3.1922 | 0.266017 | 0.07914 |
| JN087872_g | 12 | 3.8791 | 0.323258 | 0.518839 |
| *Planktophila* | 12 | 4.9744 | 0.414533 | 0.630999 |
| *Polynucleobacter* | 12 | 5.2169 | 0.434742 | 0.17564 |
| Sphingomonadaceae_uc | 12 | 5.6299 | 0.469158 | 0.239249 |
| AY532578_g | 12 | 6.8333 | 0.569442 | 0.854341 |
| *Bdellovibrio* | 12 | 4.3863 | 0.365525 | 0.130646 |
| *Cellvibrio* | 12 | 9.3697 | 0.780808 | 0.641694 |
| EU803579_g | 12 | 1.7869 | 0.148908 | 0.117079 |
| *Fluviicola* | 12 | 4.3954 | 0.366283 | 0.286912 |
| HQ343229_g | 12 | 2.384 | 0.198667 | 0.249013 |
| LBRP_g | 12 | 7.924 | 0.660333 | 0.883163 |
| LCRP_g | 12 | 2.3479 | 0.195658 | 0.094942 |
| *Nanopelagicus* | 12 | 10.4248 | 0.868733 | 4.533814 |
| Oxalobacteraceae_uc | 12 | 9.1001 | 0.758342 | 0.399625 |
| PAC000016_g | 12 | 6.8106 | 0.56755 | 0.320557 |
| Paenibacillaceae_uc | 12 | 11.7132 | 0.9761 | 11.27143 |
| *Paenibacillus* | 12 | 4.6467 | 0.387225 | 1.413813 |
| Pedobacter_g3 | 12 | 1.5937 | 0.132808 | 0.106075 |

| | | | | |
|---|---|---|---|---|
| *Sediminibacterium* | 12 | 3.6725 | 0.306042 | 0.333551 |
| *Acidovorax* | 12 | 6.9365 | 0.578042 | 0.442103 |
| *Lampropedia* | 12 | 5.2395 | 0.436625 | 0.112086 |
| *Methylomonas* | 12 | 2.5097 | 0.209142 | 0.377205 |
| *Rhizobacter* | 12 | 7.9287 | 0.660725 | 0.293389 |
| *Saccharimonas* | 12 | 3.8207 | 0.318392 | 0.116723 |
| Selenomonadaceae_uc | 12 | 2.7298 | 0.227483 | 0.133856 |
| AF370880_g | 12 | 3.6005 | 0.300042 | 0.171381 |
| *Bacteroides* | 12 | 1.5219 | 0.126825 | 0.092735 |
| CP011215_f_uc | 12 | 17.8864 | 1.490533 | 3.116723 |
| Chthoniobacteraceae_uc | 12 | 3.8789 | 0.323242 | 0.620249 |
| Comamonadaceae_uc* | 12 | 217.7156 | 18.14297 | 164.9515 |
| Cytophagaceae_uc | 12 | 5.7242 | 0.477017 | 0.859744 |
| GQ387490_g | 12 | 14.3576 | 1.196467 | 10.34723 |
| HQ827934_g | 12 | 4.6018 | 0.383483 | 0.444987 |
| Methylophilaceae_uc | 12 | 5.9172 | 0.4931 | 0.207527 |
| *Pseudarcicella* | 12 | 8.8489 | 0.737408 | 1.999995 |
| *Rheinheimera* | 12 | 11.6233 | 0.968608 | 2.543325 |
| *Sphingomonas* | 12 | 4.8219 | 0.401825 | 0.143136 |

One way-ANOVA

| *Source of Variation* | *SS* | *df* | *MS* | *F* | *P-value* | *F crit* |
|---|---|---|---|---|---|---|
| Between Groups | 4729.362 | 55 | 85.9884 | 6.738576 | 8.26E-36 | 1.354723 |
| Within Groups | 7860.541 | 616 | 12.76062 | | | |
| Total | 12589.9 | 671 | | | | |

Significant if difference in variance between two compared genera was greater than calculated LSD value (2.863933).
*Statistically significant among all other identified genera.

**Supplementary Table 5**. Short length 16S rRNA reads classified to genera level, accounting for relative abundance >1%, <1% and remains unclassified analyzed using EzBioCloud.

| Samples | Genus | | Species | |
|---|---|---|---|---|
| | ETC (<1%) | Unclassified | ETC (<1%) | Unclassified |
| R-F1-E | 22.3221 | 6.851 | 23.1975 | 50.9293 |
| R-S1-E | 45.7396 | 10.4831 | 37.9949 | 55.8858 |
| R-S2-W | 60.9992 | 9.4146 | 53.1965 | 43.7102 |
| R-F2-W | 44.5452 | 11.1655 | 33.3572 | 55.2932 |
| S-S3-N | 34.3494 | 6.4156 | 25.6981 | 33.1059 |
| S-F3-N | 19.3498 | 7.8432 | 27.5119 | 33.3124 |
| R-S4-SE | 44.1636 | 21.5543 | 41.874 | 52.4737 |
| R-F4-SE | 34.7894 | 19.0581 | 34.4392 | 62.2295 |
| R-S5-SE | 61.8705 | 11.0308 | 54.319 | 41.6809 |
| R-F5-SE | 41.0837 | 11.2373 | 32.9173 | 54.4671 |
| T-S6-N | 24.7597 | 4.8263 | 24.3198 | 11.2104 |
| R-S7-N | 36.087 | 8.9611 | 38.4933 | 20.2074 |

**Supplementary Table 6.** Oxford Nanopore MinION 16S rRNA sequencing and analyses results of sample R-F1-E and S-F3-N. The EPI2ME Fastq16S pipeline was used for the analyses.

| Samples | Sequencing replicate | Unclassified reads | No. of classified genera | No. of classified species |
|---|---|---|---|---|
| R-F1-E | Replicate 1 | 964 | 768 | 1835 |
| | Replicate 2 | 1044 | 794 | 1926 |
| S-F3-N | Replicate 1 | 717 | 714 | 1674 |
| | Replicate 2 | - | - | - |

No valid reads were obtained with second replicate of sample S-F3-N.
